# Ligands, receptors and transcription factors that mediate inter-cellular and intra-cellular communication during ovarian follicle development

**DOI:** 10.1101/594234

**Authors:** Beatriz Peñalver Bernabé, Teresa Woodruff, Linda J Broadbelt, Lonnie D Shea

## Abstract

Reliably producing a competent oocyte entails a deeper comprehension of ovarian follicle maturation, a very complex process that includes meiotic maturation of the female gamete, the oocyte, together with the mitotic divisions of the hormone-producing somatic cells. In this report, we investigate mice ovarian folliculogenesis *in vivo* using publically available time-series microarrays from primordial to antral stage follicles. Manually curated protein interaction networks were employed to identify autocrine and paracrine signaling between the oocyte and the somatic cells (granulosa and theca cells) and the oocyte and cumulus and mural cells at multiple stages of follicle development. We established protein binding interactions between expressed genes that encoded secreted factors and expressed genes that encoded cellular receptors. Some of computationally identified signaling interactions are well established, such as the paracrine signaling from the oocyte to the somatic cells through the secreted oocyte growth factor *Gdf9*; while others are novel connections in term of ovarian folliculogenesis, such as the possible paracrine connection from somatic secreted factor *Ntn3* to the oocyte receptor *Neo1*. Additionally, we identify several of the likely transcription factors that might control the dynamic transcriptome during ovarian follicle development, noting that the YAP/TAP signaling is very active *in vivo*. This novel dynamic model of signaling and regulation can be employed to generate testable hypotheses regarding follicle development, guide the improvement of culture media to enhance *in vitro* ovarian follicle maturation and possibly as novel therapeutic targets for reproductive diseases.

## INTRODUCTION

The production of a competent female germ cell line, oocyte, that can undergo fertilization requires a highly orchestrated paracrine, autocrine, endocrine and juxtracine signaling that has to occur between the oocyte and the supporting somatic cells, granulosa and theca cells. This complex biological structure formed by the oocyte and surrounding somatic cells is called ovarian follicle. During ovarian follicle maturation, a primordial follicle (50 μm diameter in the mouse) that is composed from a handful of cells, has to grow into an antral follicle (500 μm), in order to attain a competent oocyte. Attempts to separate the different cellular follicular components to study ovarian follicle maturation lead to different behavior of the individual cell types. Unfortunately, little can be controlled *in vivo* to learn the effect of different biological variables (e.g., effects of hormones, extracellular matrix stiffness) on the follicle maturation. Thus, several *in vitro* systems that mimic *in vivo* ovarian follicle development (Eppig and OBrien 1996; O’Brien et al. 2003; Xu et al. 2006) has led to some of the most significant advances in reproductive biology (Edson et al. 2009). Up to date, only organ-on-a-chip technology that combines ovarian tissue, fallopian tubes and utero(Xiao et al. 2017)— EVATAR™—has been able to mimic the ovulation period. While with EVATAR™ the effects of hormones or the extracellular matrix stiffness could be systematically study, it suffers from the same limitations as *in vivo* to understand inter-cellular and intra-cellular communications between the different ovarian follicle cell types. One of the main hindrance is the difficulty to determine how paracrine (e.g., between the oocyte and granulosa cells) and autocrine (e.g., oocyte ligands that affect the oocyte) communication, i.e., inter-cellular communication, between the different follicle cell types occurs. While some of the inter-cellular ligands such as GDF-9 and BMP-15 have been established (Knight and Glister 2006), not all of them are known. Similarly, once that a given ligand binds to its corresponding receptors, a complex signaling takes place through several biochemical mechanisms (e.g., phosphorylation, protein binding, calcium release) that ends in the activation or deactivation of transcriptional programs, i.e., intra-cellular communication. Transcription factors (TFs) are the mediators between the cytoplasmic to the nucleus signaling, by translocating between the two cellular compartments. Once that a TF is in the nucleus, directly or in form of a protein complex, it binds DNA and starts or inhibits transcription. Therefore, TFs are potent regulators of the cellular phenotype. For instance, at the follicle level, FOXL2 is a marker of granulosa cells and essential for proper ovarian follicle development(Uhlenhaut and Treier 2006).

In the recent years, advances in high-throughput techniques have allowed obtaining large amount of information about the ovarian follicle transcriptome (Chronowska 2014; Pan et al. 2005b; Skory et al. 2013; Wigglesworth et al. 2015a; Yoon et al. 2006). Analysis of these large biological datasets requires statistical and computational methods to identify the processes that are associated with the manifest phenotypes. Yet these transcriptional data have not been explored to their maximum potential. Currently, there are methods to computationally identify the more plausible TFs that are regulating a given phenotype (Grant et al. 2011; Zhao and Stormo 2011). Similarly, given a set of genes that are expressed in a given cell, the most likely genes that encode for ligands and receptors that present in a cell type could be identified, using well-curated biological databases (e.g., DIP(Xenarios et al. 2000), MetaCore). Here, we propose a system biology approach to computationally reveal the key intra- and inter-cellular dynamic processes during mice ovarian folliculogenesis *in vivo* between and among the different follicular cell types (e.g., oocyte, granulosa cells, cumulus cells) involved in each developmental stage (e.g., primordial to primary ovarian follicles).

## RESULTS

### Identification of the ligands and receptors that lead ovarian follicle development inter-cellular signaling

Understanding inter-cellular communications during ovarian follicle maturation *in vivo* entails the identification of the secreted proteins (i.e., ligands) and available receptors in oocyte and in the somatic cells (e.g., granulosa and theca cells) that support the oocyte growth and maturation. We established the most likely ligands and receptors during ovarian follicle development by characterizing the set of statistically significant genes that encode for ligands and receptors during ovarian follicle maturation. We mined several publically available time-series transcriptomics—i.e., oocytes (Pan et al. 2005a), somatic cells—e.g. granulosa and theca cells(Peñalver Bernabé et al.)—cumulus and mural cells collected during antrum formation (Wigglesworth et al. 2015b) and cumulus cells during oocyte competence acquisition (Charlier et al. 2012). Combination of all these data sets led to a list of the significant transcribed ligands and receptors in each individual cell type (e.g., oocyte, somatic, cumulus granulosa cells).

We identified 100 genes that encode for ligands and 95 genes that encode for receptors that could potentially regulate the inter-cellular communication during ovarian follicle development (**File S1**). Some of the genes that encode for ligands and receptors were active in multiple cell lines, e.g., *Dnc* in the oocyte and somatic cells, *Efna2* in mural and cumulus granulosa cells; yet others were very specific (e.g., *Wnt10a* in cumulus cells). More than half of the genes that encode for ligands, a total of 59, were cell-specific: 12 to the oocyte, 19 in somatic cells, 8 in cumulus granulosa cells, 10 mural granulosa cells, and 10 in cumulus granulosa cells during the oocyte transition from a chromatin non-surrounded nucleolus (NSN) to a surrounded chromatic nucleolus (SN). More than 60% of the receptors were specific (i.e., 12, 24, 3, 12, and 7 in the oocyte and somatic, cumulus and mural granulosa cells and in cumulus cells during the oocyte transition from NSN to SN, respectively). In terms of the number of stages that a given gene that encode for ligand or a receptor was active, some intercellular signaling proteins were more ubiquitous as they were active at multiple ovarian follicle stages (e.g. *Apoa4* from secondary to large antral follicles, *Igf1* from primary to large antral follicles), while others were very specific, such as *Bmp15* and *Wnt6*, which were only active during the primordial to primary transition or in the primary to secondary transition, respectively. Finally, we identified several genes that encode for ligands that have been reported to bind to multiple genes that encode for receptors, e.g., *Thbs1* or *Vegfa* (55 and 37 connections during the small to large antral transition, respectively). Yet some were very specific as only bind 1 or 2 receptors, e.g., *Shh* or *Rspo2* (**File S1**).

### Constructions of inter-cellular signaling networks during ovarian follicle maturation

Combination of multiple datasets and manually curated databases—i.e., Metacore and DIP(Xenarios et al. 2000)—led to the identification of a total of 1,663 connections between the 100 ligands and 95 receptors (**File S1**). Out of those interactions, 46% of the connections were autocrine signaling, mostly between somatic cells (45%)—note that these somatic autocrine connections could be within the same somatic cell or between two different somatic cells within the same follicle or even between two different follicles. In terms of the possible paracrine signaling, more than 22% were initiated from a ligand produced by the oocyte. Interestingly, 290 of the 1,663 connections that we identified were between specific genes that encode for ligands and receptor (i.e., only significant in one cell type) and 40% of them were autocrine signaling, mostly within somatic cells (65 somatic ligands to somatic receptors).

### Inter-cellular signaling networks *in vivo from primordial to primary*

The primordial to primary transition was the most complex of all the stages during ovarian follicle development. The majority of inter-cellular communications occurred during the transition from primordial to primary ovarian follicles (22% of the connections). Multiple hallmarks during this transition were present in the transcriptomics data (e.g., zona pellucida formation, gap connections) and multiple genes known to be involved in the development of primary ovarian follicles from primordial ovarian follicles were identified in the transcriptional data (e.g., *Zp1, Gja4, Amhr*, **Supplemental Note 1**).

The transcriptional activity and the number of paracrine and autocrine communications of the somatic cells surpassed those of the oocyte (**Table S1**). Out of all the autocrine and paracrine communications, only a few of them were cell specific, i.e., between ligands and receptors predicted as uniquely present in one cell type (**File S1**). Precisely, only 11 oocyte autocrine, 15 somatic autocrine and 12 oocyte-somatic and 28 somatic-oocyte paracrine communications were specific. Our results recovered well-known ligands in this stage, such as *Gdf9, Bmp15* or *Amh* as well as ligands that have not been previously reported in the literature of their presence and role during the primordial to primary ovarian follicle transition, such as *Adam2* and *Ntn3* (**Fig. 2**)

The intricacy and complexity of the primordial to primary transition is clearly depicted in the inter-cellular networks, which were dived into several subnetworks. The largest subnetwork (shown in green in **Fig. 1**) included well-known ligands and receptors from the *Tgf* family (e.g., *Gdf9, Bmp4, Inha*) and from the *Bmp* family and also contained a substantial core of diverse extracellular binding protein families, such as integrins (e.g., *Itga6, Itgb1*), laminin (e.g., *Lama1, Lamab1*) and collagen (e.g., *Col18a1*). Only the integrins and laminins transcribed in the oocyte (i.e., *Itga5, Itgb1, Lamb1*) significantly changed in this transition (p-value<0.01, **File S1**). Some ligands, such as *Dcn*—proteoglycan identified in the later stages of follicle development (Adam et al. 2012)—and *Nrp1* were produced by both the oocyte and the somatic cells. Additionally, this green subnetwork contained connections not previously study in relationship to ovarian follicle development, such as *Ntn3* and their corresponding receptors, e.g., *Unc5b, Unc5c* and *Neo1*. According to our computational model, *Ntn3* was a somatically-produced ligand that interacted with receptors in oocyte and the somatic cells. *Ntn3* functions have been described in other developmental processes, such as axonal growth (Kang et al. 2004; Wang et al. 1999). Similarly, its receptors *Unc5b* and *Unc5* are known to participate in angiogenesis(Larrivee et al. 2007; Lu et al. 2004) and are anti-apoptotic (Ozmadenci et al. 2015) and *Neo1* is related with cellular growth(Wilson and Key 2007).

**Figure 1.**
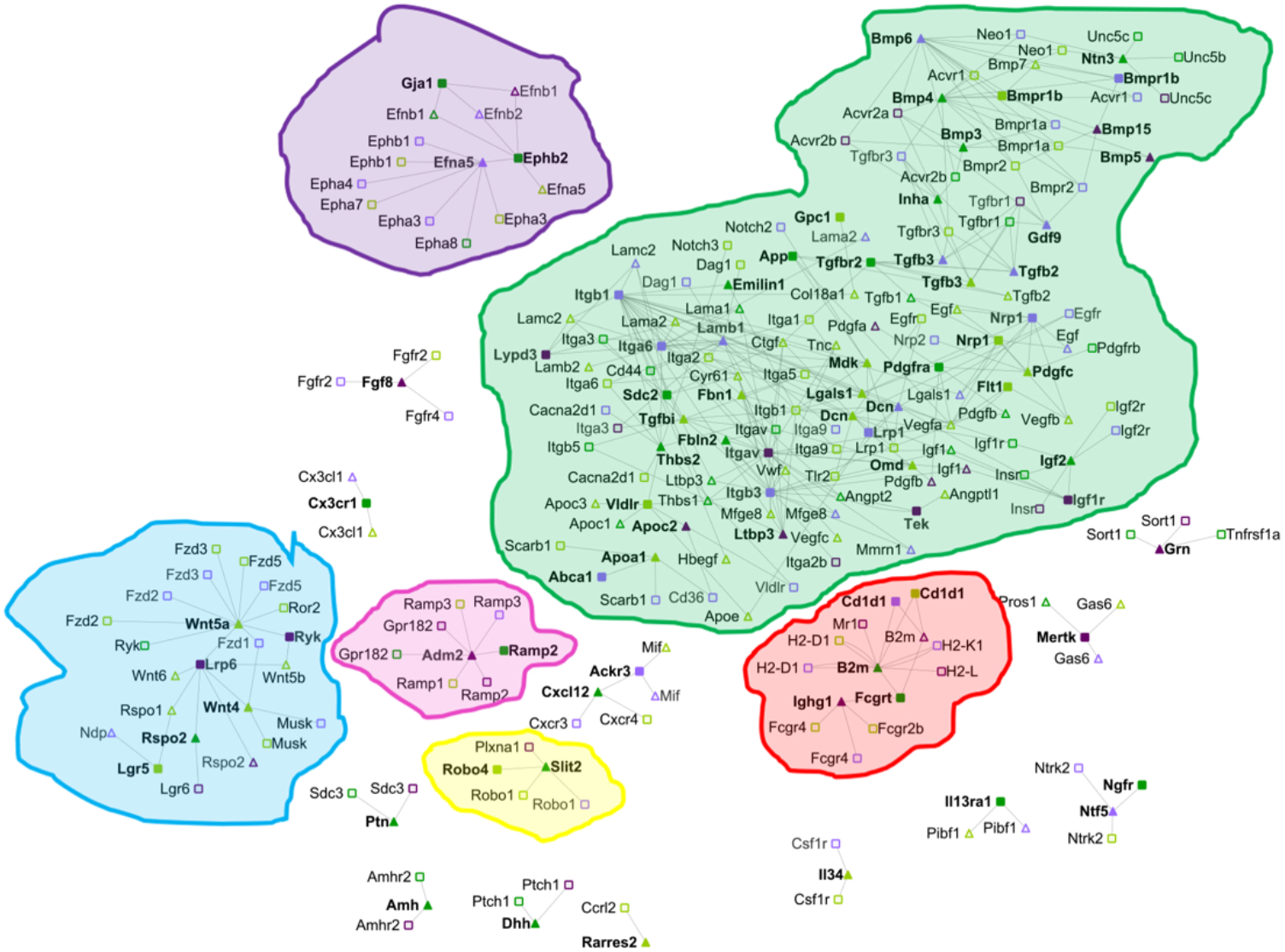
Intercellular networks between oocytes and somatic cells primordial to primary transition. Receptors only identified as significant (FC≥2 and p-value≤0.01) in the somatic cells are represented as darker green squares (e.g., *App, Tgfbr2*); ligands only identified as significant (FC≥2 and p-value≤0.01) in the somatic cells are represented as darker green triangles (e.g., *Amh, Thbs2*); Receptors only identified as significant (FC≥2.5 and p-value≤0.01) in the oocyte are represented as darker purple squares (e.g., *Itga9, Bmpr1b*); ligands only identified as significant (FC≥2.5 and p-value≤0.01) in the oocyte are represented as darker purple triangles (e.g., *Tac1, Bmp5*). Lighter colors indicate that the receptor or ligand are also significant in other cell types during ovarian follicle development (e.g., *Nrp1, Tgfb3, Bmpr1a, Gdf9*). Bold text corresponds to genes whose abundance change during the specific transition (e.g., primordial to primary), such as *Gdf9, Neo1*; otherwise, those genes are presented in the follicles during the given transition yet their abundance is not changing (e.g., *Egfr, Sdc3*) are not bolded. Connections between the different ligand-receptors are only present if at least one of member of the pair is changing its expression during a given developmental stage. The connections and the references for each of the edges in the network can be found in **File S1**.

**Figure 2.**
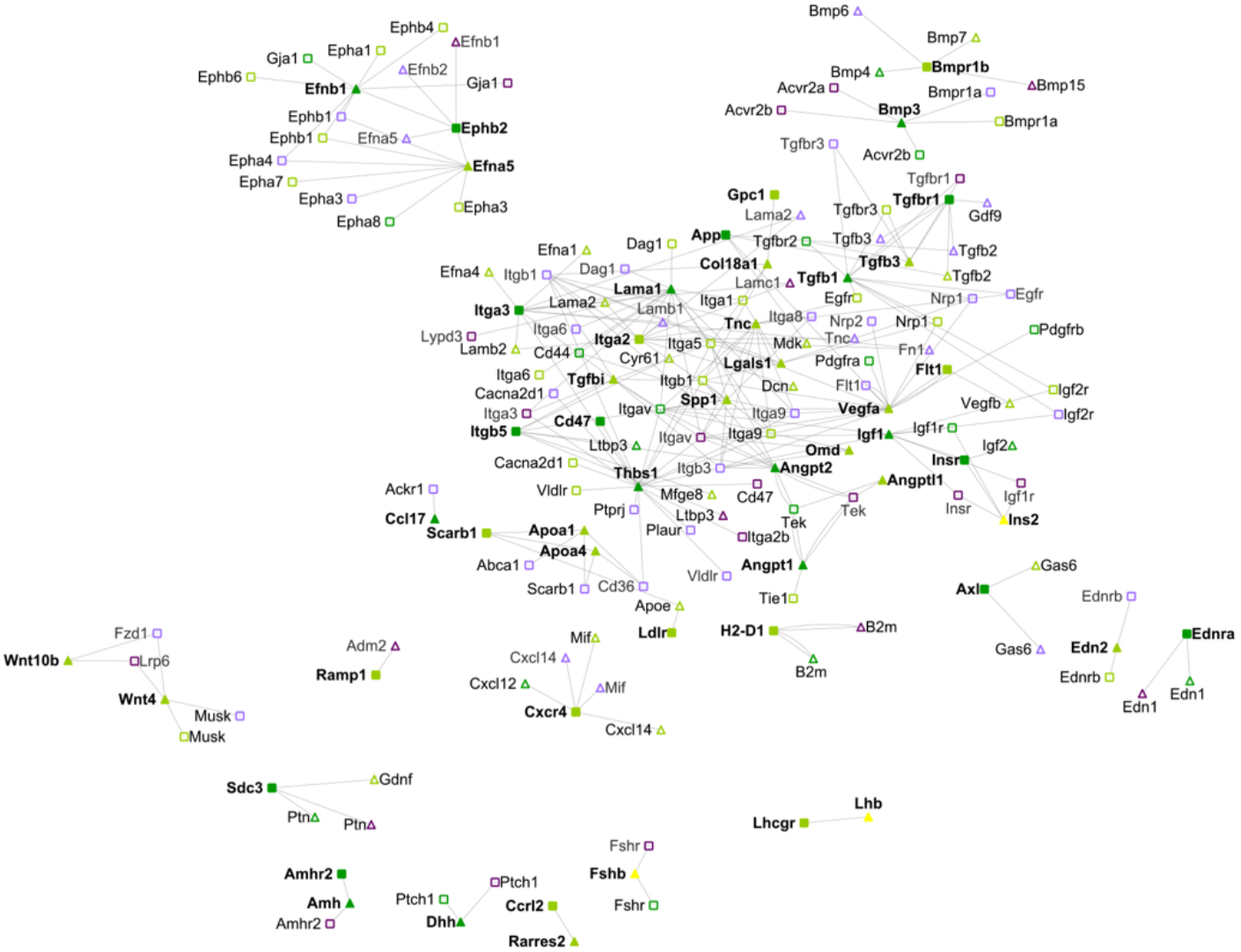
Intercellular networks between oocytes and somatic cells from small antral to large antral transition. Legend as in **Fig. 2**. Yellow nodes indicate ligands secreted by the endocrine systems and present in the blood accessible to the growing antral follicle (e.g., *Lhb, Ins2, Fsh*). The connections and the references for each of them can be found in **File S1**.

Other medium size networks (e.g., purple) encoded the *Ephrin* and *Wnt* families. Oocyte secreted *Efna5* ligand interacted with *Epha1* and *Gji* somatic receptors, which were among the top genes whose transcriptional abundance changed the most (**File S1**). Mice who lack *Efna5* ligand are subfertile (Buensuceso et al. 2016). Also, this purple subnetwork encompassed paracrine communication from somatic ligands *Wnt4* and *Wnt5* to oocyte specific receptors *Lrp6* and *Ryk. Wnt4* signaling regulate the expression of *Amh* and mice that lack Wnt4 suffer from premature ovarian failure(Prunskaite-Hyyrylainen et al. 2014). *Rspo2* somatic specific ligand also bind to the oocyte specific receptor *Lrp6*. The red highlighted subnetwork contained a somatic autocrine communication between *B2m* and *Fcgrt* and the later was also involved in a specific oocyte-somatic communication with the oocyte ligand *Ighg1. B2m* and *Igh1* have been previously studied in relation to ovarian cancer biology(Qian et al. 2018; Yang et al. 2009), but this research has not been extended to ovarian follicle development.

Finally, two other smaller and disconnected networks were related with the *Ramp* and the *Robo* families of receptors. *Ramp2* was identified as a specific somatic receptor that could bind to the oocyte specific ligand *Adm2*—pink subnetwork. *Adm2* prevents oocyte atresia by regulating cell-cell interactions in cumulus-oocyte complexes (Chang et al. 2011). The specific somatic receptor *Silt2* could bind to the non-specific somatic ligand *Robo4*—yellow subnetwork. This interaction between *Silt2* and *Robo* is known to occur at the time of the formations of the ovarian follicles and diminished the rate of oocyte proliferation(Dickinson et al. 2010). Finally, neurotrophins soluble growth factors were also identified as important during the primordial to primary transition, corroborating previous studies (Dissen et al. 1995) and their involvement on the formation of squamous somatic cells (Dissen et al. 2001) through the oocyte-somatic specific paracrine communication between *Ntf5* and *Ngfr*.

### Inter-cellular signaling networks *in vivo from primary to secondary*

During the transition between primary to secondary follicles, the follicle start acquiring up to 10 layers of granulosa cells (Pedersen and Peters 1968), the formation of the theca layer commences and the follicles are capable to produce estrogenic hormones. Several of the transcripts involved in those developmental pathways could be recapitulated from the recompiled ensemble transcriptional data, such as *Cyp17a1* (**Supplemental Note 2**). Numerous transcripts were changing during the primary to secondary transition, most likely due to the addition of the theca cells, yet the complexity of the inter-cellular signaling network was reduced compared with the primordial to primary transition (**Table S1, Fig. S1**). The majority of the intercellular communications during the primary to secondary transition were autocrine communications between somatic cells, followed by paracrine signaling between ligands secreted by the somatic cells to receptors in the oocyte. Only 21 of the inter-cellular communications were specific, i.e., there were between ligands or receptors only expressed in the oocyte or somatic cells. For instance, *Angpt2* was produced by the somatic cells and interacted with integrins *Itga5* and *Itgb5* that were present only in somatic cells and with an oocyte specific receptor, *Tek. Angpt2, Igf2, Pros1*, and *Thbs2* were the only cell-specific genes that encode for ligands and *H2-D1, H2-L, Tgfbr2* and *Sdc3* the only transcripts encoding for cell-specific receptors that were significantly altered during the primary to secondary transition.

In terms of sub-networks, communities highlighted in green, pink, and blue during the primordial to primary transition (**Fig. 1**) were diminished in terms of the number of connections between the transcripts and the yellow and purple communities were not present at all (**Fig. S1**). Similarly, the somatic-oocyte paracrine and somatic autocrine communications of *Amh* were not identified either. Interestingly, a somatic-specific subnetwork of the *Edn* family appeared during the primary to the secondary transition, in agreement with prior studies of the role of endothelin in ovarian follicle development(Bridges et al. 2011)

### Inter-cellular signaling networks *in vivo* from secondary to small antral transition

Only a few secondary follicles sensitive to endocrinal hormones FSH and LH will transition into small antral follicles, avoiding atresia, the default pathway (McGee and Hsueh 2000). Follicles start producing androgens in the theca cells and estrogens in the granulosa cells (Wood and Strauss 2002), the antrum cavity emerges, filled with hyaluronic acid and proteoglycans(Gebauer et al. 1978; Jensen and Zachariae 1958), such as versican and perlican (Eriksen et al. 1999), and theca cells become vascularized(Young and McNeilly 2010). Multiple of the genes that are known to play a role during this transition were also significantly changing, e.g., *Fshr, Vcan* (**File S2, Supplementary Note 3**), although the number of downregulated genes exceeded the number of upregulated transcripts (**Table S1**). The complexity of the inter-cellular signaling network during the secondary to small antral transition was similar to the primary to secondary transitions, and thus with less inter-cellular connections than the primordial to primary transition (**Fig. S2, Table S1, File S1**). The majority of inter-cellular communications were somatic autocrine interactions and only 12 of them were specific. Out of the active ligands, *Inha* was the only one whose transcriptional abundance increased in this stage; *Rspo2* and *Wnt9* were significantly downregulated (**File S2**).

There were distinct changes during the secondary to small antral transition compared with the two other prior transitions. For instance, the major blue subnetwork during the primordial to primary transition (**Fig. 1**) was divided into two smaller subnetworks, one of them highly enriched in members of the *Tgf* family (**Fig. S2**). Somatic cells started been sensitive to INS2 and FSH and *Wnt* signaling intercellular communications were more prevalent that in the primary to secondary transition. While the *Ephrin* family networks appeared again at this stage, the *Edn* subnetwork, important in the primary to secondary transition disappeared.

### Inter-cellular signaling networks *in vivo* from small to large antral transition

At the end of this transition, the oocyte is competent to resume meiosis (Mehlmann 2005) and the large antral cavity that allow enough oxygen supply to the oocyte is fully formed. Transcripts from genes that are known to participate during the antrum formation were present in the transcriptomic data that we collected (e.g., *Hspg2, Star, Hsd3b1*, **Supplemental Note 4, File S2**). The oocyte was mostly transcriptionally silent, yet the somatic cells were very active—even more so than in any other prior stage during ovarian follicle development, which the number of downregulated genes excessed the number of upregulated transcripts (**Table S1**). Opposite to the two prior transitions—from primary to small antral follicles—the complexity of the signaling network highly raised and communications were led by somatic cells (**Fig. 2**). Indeed, all the autocrine communications emerged between somatic cells and the majority of the paracrine signaling was through somatically-produced ligands. Only 5 inter-cellular communications were between actively changed and cell specific transcripts (i.e., oocyte or somatic cells) and all of them were somatic autocrine connections (e.g., *Col18a1* and *Gpc1*).

The large and complex green subnetwork that involved members of the *Tgf* family, integrins and vascular signals during the primordial to primary transition (**Fig. 1**) appeared again during the formation of the antral cavity (**Fig.2**). The *Eph* family subnetwork (highlighted in purple in **Fig.1**) contained more nodes and more connections at this stage compared to the primordial to the primary transition. On the other hand, the blue subnetwork—mostly enriched in *Wnt* genes that encode for ligands—and the red subnetwork—associated with *Ramp* genes that encode for receptors—decreased their importance during the antral cavity formation and the subnetwork associated with the *Robo* family (yellow subnetwork) had completely disappeared. The connections pertaining to the *Edn* families were significant again at this stage, as they were during the primary to secondary transition (**Fig. S1**), and the transcriptional levels of the gene that encode for the *Lhcgr* receptor were significantly increasing for the first time during ovarian follicle development. Also, other endocrinal led communications, such as from Fsh to Fshr or from Ins2 to its somatic receptors, were present at this stage as well.

### Inter-cellular signaling networks *in vivo* from small antral to large antral between oocyte and mural and cumulus granulosa cells

At the phenotypical level, one of the most important biological processes during antral formation is the differentiation of the granulosa cells into mural and cumulus granulosa cells (Mehlmann 2005). Several genes involve in this stage were presented in the publically available transcriptomic data of mural and cumulus cells(Wigglesworth et al. 2015b), such as *Cd34* and *Has2* (**Supplemental Note 5, File S2**). The number of downregulated and upregulated genes was very comparable in cumulus granulosa cells and in mural granulosa cells (**Table S1**). Interestingly, the number of total transcripts that were significantly changed in the cumulus cells far exceeded those of the oocyte, somatic cells or mural granulosa cells (**Table S1**). More than a third of the significantly altered genes in the cumulus cell transcriptomic data were specifically produced bynonly cumulus granulosa cells. The transcripts from mural granulosa cells exhibited a similar ratio of specificity.

The number of paracrine and autocrine signals was substantial, with almost all the autocrine signaling equally divided between mural or cumulus granulosa cells (**Table S1, Fig. 3**). Several paracrine communications were initiated by non-significantly changing oocyte ligands to receptors in both mural and cumulus cells and the order of magnitude of paracrine communications for cumulus and mural granulosa cells were comparable, with a limited number towards non-significantly changing receptors in the oocyte.

**Figure 3.**
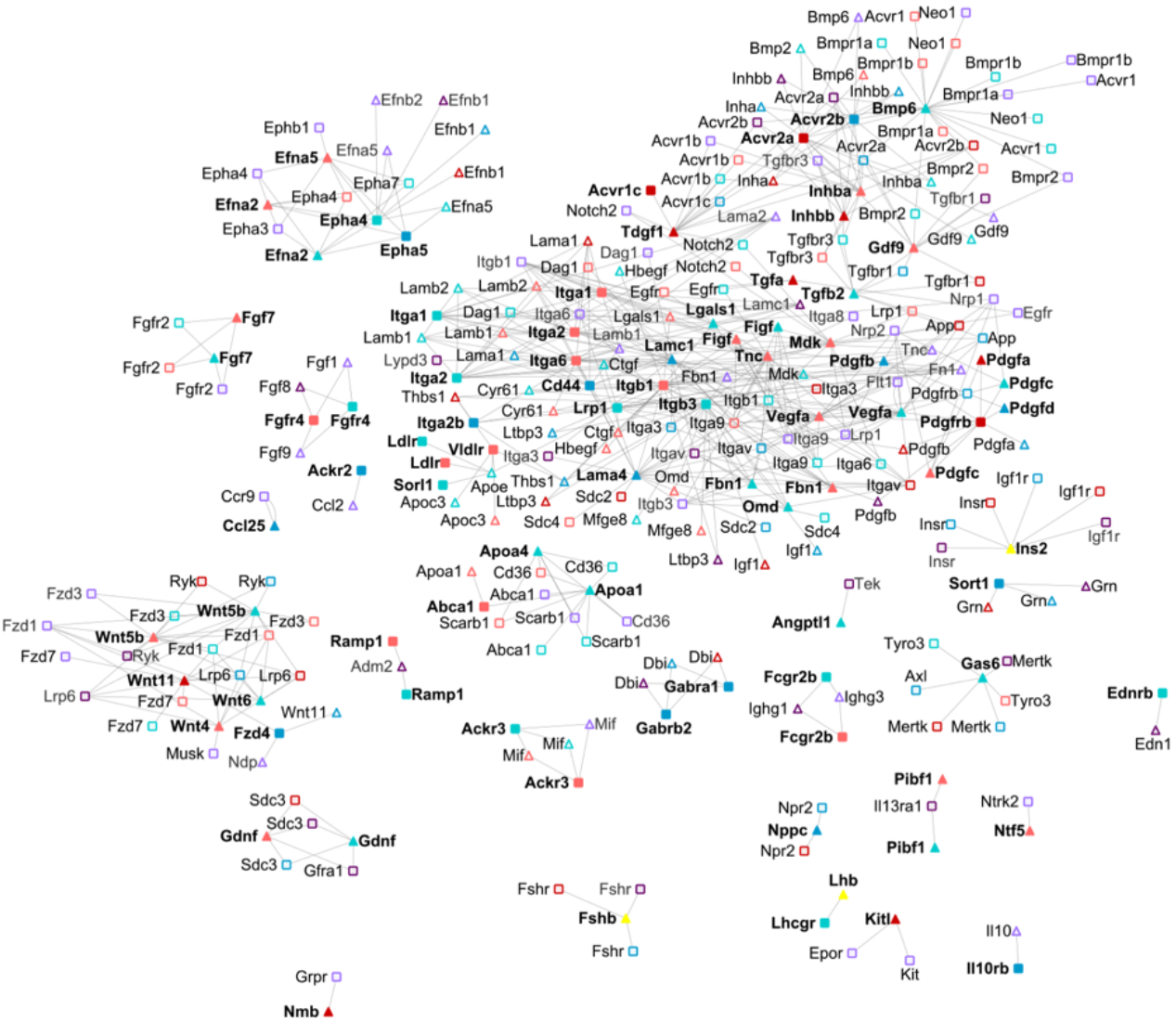
Intercellular networks between oocytes and cumulus and mural cells from small antral to large antral transition. Legend as in **Fig. 2**. Red, cumulus cells; blue, mural cells; yellow ligands from the endocrine system. The connections and the references for each of them can be found in File S1. Oocyte: FC≥2.5, p-value≤0.01; cumulus and mural granulosa cells: FC≥2 and p-value≤0.01. The connections and the references for each of them can be found in **File S1**.

The network inter-cellular signaling pathways specificity seemed to agree with the distinct and specific functionality that mural and cumulus granulosa cells play during ovarian follicle development. For instance, while *Gdf9* transcript was detected in the cumulus, mural and oocyte, it was only actively changing in cumulus cells. Several ligands (e.g., *Wnt11, Tgfa, Inhba*) and receptors (e.g., *Pdgfrb, Acvr1c* and *Acvr2a*) were specifically produced by the cumulus cells, while ligands such *Epha5, Garba1, Sort1* and *Pfgfd* and receptors such as *Fzd4, Il10rb, Cd44* or *Gabrb2* were specific to mural cells. The autocrine signaling between the ligand *Pdgf* and the receptor *Pdgfrb* only occurred in cumulus cells and the paracrine communication between cumulus ligand *Wnt11* to mural receptor *Fzd4* was specific as well. At least at the transcriptional level, the mural autocrine communication between the ligand *Lamc1* and the receptor *Cd44* was important, with abrupt transcriptional increases for both genes (FC=265, p-value (fdr)=5.48*10^-17^ and FC=135, p-value (fdr)=8.68*10^-18^, respectively).

At the subnetwork level, the green subnetwork during the primordial to primary transition (**Fig. 1**) gained new members of the *Tgf, Pdgf* and integrin families. More members of the *Apoa* family were detected, although fragmented from the large green subnetwork (**Fig. 3**). Interestingly, more members of the *Fgf* family emerged as well as multiple small subnetworks related with *Kitl, Il10, Il13ra1* and *Gdnf*. The *Wnt* family become mostly focused in *Fdz* receptors and also gained more connections. On the contrary, *Amh* signaling completely vanished at this mural-cumulus-oocyte network stage (**Fig. 3**).

### Inter-cellular signaling networks *in vivo* during the NSN to SN transition in the oocyte

Oocyte maturation is required for adequate egg fertilization. One of the hallmarks required for achieving oocyte maturation is chromatin condensation (Albertini et al. 2003), i.e., chromatin condensation in the oocyte nucleolus, from a non-surrounded nucleolus (NSN) to a surrounded nucleolus (SN) stage (Zuccotti et al. 1995). This transition cannot be achieved by nude oocytes; they required the present of cumulus granulosa cells (De la Fuente and Eppig 2001). Analysis of the available transcriptomics data during the NSN to SN transition agreed with the current understating of the changes associated with oocyte maturation (**Supplemental Note 6**).

Cumulus cells were orchestrating the NSN to SN transition with a total of 237 inter-cellular signaling communications, out of which 162 were autocrine communications between cumulus cells, 55 were paracrine communications led by cumulus cell ligands. There was no autocrine communication between the oocyte and no paracrine communication led by ligands secreted by the oocyte. While multiple genes that encode for cumulus ligands increased their transcription rates during this stage (e.g., *Wnt10a, Ltb, Il6, Ereg, Camp* and *Apln*), their associated receptors did not significantly change their transcription levels (i.e., *Aplnr, Cd14, Erbb2, Erbb4, Il6ra, Il6st, Ltbr, Robo1, Robo2, Fzd1* and *Lrp5*). Additionally, *Gpr182* and *Epha8*, cumulus cell specific genes according to our models, were significantly upregulated in cumulus cells. As expected, the oocyte was completely silent at the transcriptional level (**File S2**).

At a more granular level, the large subnetwork during the primordial to primary transition (marked in blue in **Fig. 1**) contained less nodes and less interactions during the NSN to SN transition (**Fig. S3**). For instance, the cumulus specific ligand *Fd6* and its associated connections were non-present. Similarly, *Serpine1, Ins2, Erg*, and *Tgf* families were separated from the majority of the components of the core blue subnetwork, which still contained a large number of inter-cellular communications through integrins (e.g. *Itga7*). Other parts of the primordial-to-primary blue subnetwork completely vanishes, such as *Gdf9, Bmp and Inhibin* families. Additionally, multiple small subnetworks only appeared during this stage (i.e., *Ifnr1, Cd47* and *Gpr182*; *Il6; Camp)*; while others disappeared (e.g., *Ramp1, Akrc* receptor and *Fgf* ligand families). Interesting, *Robo*, which was only present during primordial to primary transition (**Fig. 1**), became significant again during NSN to SN transition.

### Identification of the most likely TFs that control the significant genes in each follicular cell type during *in vivo* follicle maturation

Finally, we identified the most likely TFs that could regulate the significant genes during ovarian follicle development *in vivo* for each cell type and each follicular stage (**Fig. 4, File S3**) by determining the targets for a given TFs from experimental, manually curated databases, e.g., Metacore, Ovarian kaleidoscope (Leo et al. 2000), or computationally, e.g., FIMO (Grant et al. 2011) and BEEML (Zhao and Stormo 2011).

**Figure 4.**
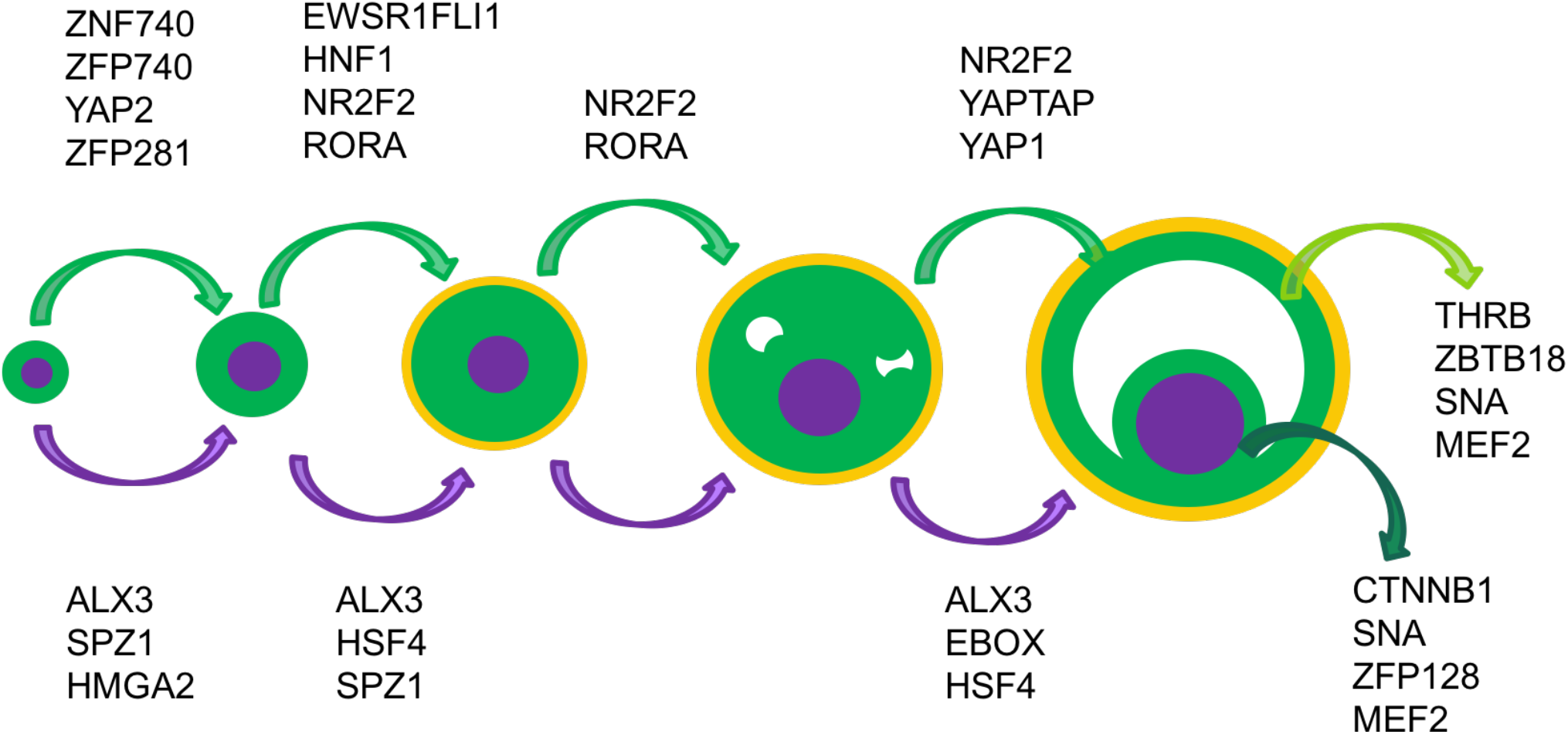
Representative active TFs during ovarian follicle development in the oocyte, somatic cells, cumulus and mural cells. The rest of the transcription factors are summarized in **File S3**.

Based on our results, TF ALX3 regulated oocyte transcriptional program from primordial to antral follicles, with the exception of secondary to small antral transition—most likely due to the lack of power from the close similarity between secondary follicles and small antral follicles. Other TFs that regulated oocyte development included SPZ1, from primordial to the secondary follicles and HSF4 from primary to antral follicles. Some TFs were specific to ovarian follicle transitions. For instance, TF MHGA2 was significant during the primordial to primary transition, while the TF EBOX was identified as a very likely regulator during the last ovarian follicle stage, from small antral to large antral transition.

For somatic cells, several zinc finger TFs (e.g., ZFP281 and ZFP740) regulated the initial activation of primordial follicles, including YAP signaling—also identified as a possible activated TF during the small antral to large antral transition. NR2F2 and RORA—RORA is a theca TF(Young and McNeilly 2010)—were among the most significant TFs from the primordial to the small antral transition and NR2F2 regulated the transcription of a large group of genes during antral formation. While some TFs were common in the mural and cumulus cells (e.g., SNA and MEF2), others were uniquely active either in cumulus cells (e.g., CTNNB1 and ZFP128) or in mural cells (e.g., THRB and ZBTB18), as expected due to the different functions that these two cells types have during ovarian follicle development.

## DISCUSSION

Signals that control ovarian follicle development and enable the formation of a competent oocyte are not fully understood yet due to the difficulty of disentangled the intra-cellular and inter-cellular communications among and between the oocyte and its surrounding somatic cells. In this report, we have presented the most plausible inter-cellular communication networks during *in vivo* follicle maturation, as well as the most likely TFs that controlled and regulated ovarian follicle development at each follicular state and in each individual follicle cell using available transcriptomic data. These inter-cellular networks and intra-cellular regulation can help to generate testable hypothesis in the laboratory that can enable to better understand this complex system.

Not only our computational approach has been able to recapitulate the presence of well-known ligands and receptors including the cell type that produce them (e.g., oocyte, mural cells) and the exact ovarian follicle developmental stage (e.g., from primordial to primary stage), but we also identified novel ligands and receptors that were non-known to play a role during ovarian follicle development. For instance, several families of ligands and receptors that are well-known to intervene during follicle development such as the *Bmp, Inh*, and *Tgfb* families and their corresponding receptors, the *Bmpr, Acvr* and *Tgfbr*, as well as mechano-transduction receptors such as integrins were included in our inter-cellular signaling networks. Yet, our transition specific networks also portrait other families that have been little explored during ovarian follicle development. For instance, the role of the *Efn* family and its receptors *Eph* (Buensuceso and Deroo 2013), or the functions associated with the binding of the *Thbs* family (Hatzirodos et al. 2014) to integrins present in the oocyte plasma membrane, especially during the primordial to the primary transition. Additionally, there are several unique genes such as *Gnr, Ighg1, Ndp, Ntn3, Pibf1, Pros1, Sct, App, Neo1, Tyro3, Ptprz1* or *Phtr1*, just to name a few, which have been barely examined in their connection to ovarian follicle development. Moreover, the combinatorial possibilities are enormous: there are ligands capable to bind to the same receptor in the oocyte and somatic cells such as *Amh*, while others bind to multiple receptors such as *Vegfa*; there are ligands whose genes are expressed in both cell types and have receptors in both cell types as well, such as *Dnc*. All this complex inter-cellular communication supports previous observations related with the difficulty to grow primordial follicles in 3D alginate gels, while it is possible to growth them in ovarian tissue(Eppig and OBrien 1996; O’Brien et al. 2003) or groups of primordial follicles(Hornick et al. 2013), as the somatic and ovarian cells proportionate all those ligands.

In line with the experimental observations of the difficulties of growing primordial follicles in vitro by themselves, the most entangled communication between the different cell types occurs between primordial to primary states through ligands that were mostly secreted from somatic cells. While the oocyte autocrine and oocyte-somatic paracrine communication proportions decreased during follicle maturation, from 64 to 0; somatic autocrine and somatic-oocyte paracrine communications, were maintained or slightly decreased (**Table S1, Fig. S4**). These results highlight the growing importance of somatic cells in controlling the intercellular communications between the oocyte and the somatic cells as the follicle matures from the primordial to the antral stage. Additionally, our inter-cellular networks between the primordial to primary transition indicated that this early stage may entitle several very convoluted communications between the oocyte and the surrounding somatic cells or very likely the stromal cells in the ovary as primordial follicles can grow in *ex-vivo* ovaries. Importantly, from the inter-cellular networks, there are several possible candidates that maybe further explored experimentally by adding them to the follicle culture media to grow *in vitro* primordial follicles, such as *Ntn3* or *Omd*.

Finally, using available experimental data and computational methods, we were able to also recovered TFs that have been studied before during ovarian follicle maturation as well as others not that well understood, such as zinc finger proteins. Our results indicate that the YAP/TAP pathway is indeed active in somatic cells (i.e., granulosa and theca cells) in the primordial to primary transition and during the granulosa cell expansion (**Fig. 4**). YAP/TAP signature is an indication of cell proliferation and growth (Lei et al. 2008), which correlates with the great cellular expansion that the granulosa cells undergo during ovarian follicle development. Yet, primordial follicles are under arrested growth due to the activation of the Hippo pathway, which in turn represses YAP/TAP activation(Kawamura et al. 2013b). In fact, the YAP/TAP pathway is activated spontaneously when arrested primordial follicles are removed from their ovaries (Kawamura et al. 2013a). While it is not clear how YAP/TAP regulation could overcome Hippo repression, one plausible mechanism is through *Akt* dephosphorization of YAP in activation in primordial follicles (Li et al. 2010) and through a FSH-mediated PKA activation in secondary follicles (Yu et al. 2013). The final size of antral follicle might be regulated by the activation of the Hippo pathway. Expansion of the granulosa cells increases the number of granulosa cell-cell interactions (Aplin et al. 1999) and mechanical stress is capable to activate the Hippo pathway (Gerard and Goldbeter 2014), thus repressing the YAP/TAP activation and subsequently avoiding the continuation of the granulosa cell proliferation.

While our computational approach is very powerful to disentangle the complex inter-cellular communications during ovarian follicle maturation, several pitfalls should be noted. For instance, though the inter-cellular connections apparently only affect a given follicle, secreted molecules, either proteins or metabolites by the oocyte and somatic cells are capable to alter the processes in other surrounding follicles as well, directly (i.e., by binding in the receptors active in other somatic cells) or indirectly (i.e., by altering the endocrine system). These differences cannot be unraveled with the current available experimental data. Moreover, as the transcriptomics data belong to surviving follicles, all the competition effects that lead some secondary follicles undergo atresia and not entering the recruiting pool and subsequently are not represented in the inter-cellular networks depicted in this article. Finally, the current networks are based on transcriptional abundance—we have employed RNA levels as a proxy of protein activation—and they can be highly improved by obtaining proteomic measurements at each follicular transition for each individual cell type (e.g. oocyte, granulosa, theca, mural and cumulus cells).

In summary, systems reproductive biology approaches not only allow to reveal the key ligands and receptors associated with each cell type at each transition during ovarian follicle development, but also understand the complex autocrine and paracrine communications between the oocyte and surrounding supporting cells that allow the production of a competent oocyte. We expect that our computational predictions allow to generate novel data based hypothesis that could be experimentally validated to increase our comprehension of ovarian follicle development and thus the exploration of novel treatments for fertility disorders, such as polycystic ovarian syndrome or fertility preservation.

## ACKNOWLEDGMENT

We thank Dr. Lei Lei, Sarah Kiesewetter for their invaluable technical support and Dr. Nadereh Jafari, Director of the Genomics Core Facility (Center for Genetic Medicine, Northwestern University) and Dr. Simon Lin and Dr. Gang Feng.

## Author contributions

BPB designed the computational approached and performed all the computational study. BPB, TKW, LJB and LDS interpreted the results and wrote the manuscript.

## METHODS

### Inter-cellular networks

Murine genes that encode for secreted proteins and receptors were identified using the GeneGO database (Advance Search 2.0). Protein-protein interactions between secreted proteins and receptors were obtained from manually curated databases, GENEGO and DIP(Xenarios et al. 2000). Autocrine and paracrine connections were deemed possible if either one of the member of the interaction, either the ligand or the receptor, has a statistically significant change for the corresponding transition under consideration—see **Table S3** from Peñalver Bernabé and colleagues for more details—and that the corresponding receptor or ligand was at least present in the microarray for the very same transition. The specificity of each genes that encode for a ligand or a receptor was previously identified—i.e., a gene is only transcribed by the oocyte, by the mural cells(Peñalver Bernabé et al.). All the inter-cellular graphs were plotted with Cytoscape (Shannon et al. 2003).

### Most likely transcription factors

Computationally predicted targets of TFs were detected by exploring whether the TF position weighted matrices (PWMs) could bind to the consensus mammalian promoter regions of a given gene (Xie et al. 2005) between −2000 to 2000 base pairs with respect to the transcription starting site (TSS) of the given gene. We used to different TF binding site search programs, FIMO (Grant et al. 2011) and BEEML (Zhao and Stormo 2011), to establish the targets of a TF. Agreement between the results of FIMO and BEEML, using cutoff of p-value≤10-4 and E-score≥0.3, respectively, was deemed as an indication that a given TF could bind to the promoter region of a gene. The list of explored TFs using FIMO and BEEML was obtained from a combination of different sources: i) TFs that have experimentally identified position weighted matrices (PWMs) in vertebrates that were reported in TRANSFAC(Matys et al. 2003; Wingender et al. 1996); ii) the non-overlapping PWMs that were in the CIS-BP database (Weirauch et al. 2014) but not in the TRANSFAC, including the non-inferred PWMs in the CIS-BP dataset –a total 3,216 PWMs from 1,164 different TFs. We additionally added connections between TFs and target genes that were in the Ovarian kaleidoscope database (Leo et al. 2000), and from conserved motifs in mammals (Schulz et al. 2012). We established whether a TF was active at a given stage by determining the significance of the ratio between genes that were significantly changing for a given cell type and stage compared with non-significant genes using a hypergeometric distribution (Sui et al. 2005). A total of 500 bootstrapping samples with the same number of the non-significant genes for a given cell type and stage than the number of significant genes were selected. Medians for p-values were reported after been corrected for multiple comparisons using false discovery rate method (Benjamini and Hochberg 1995).

## SUPPLEMENTARY FigURES, TABLES, NOTES AND FILES

**Figure S1.**
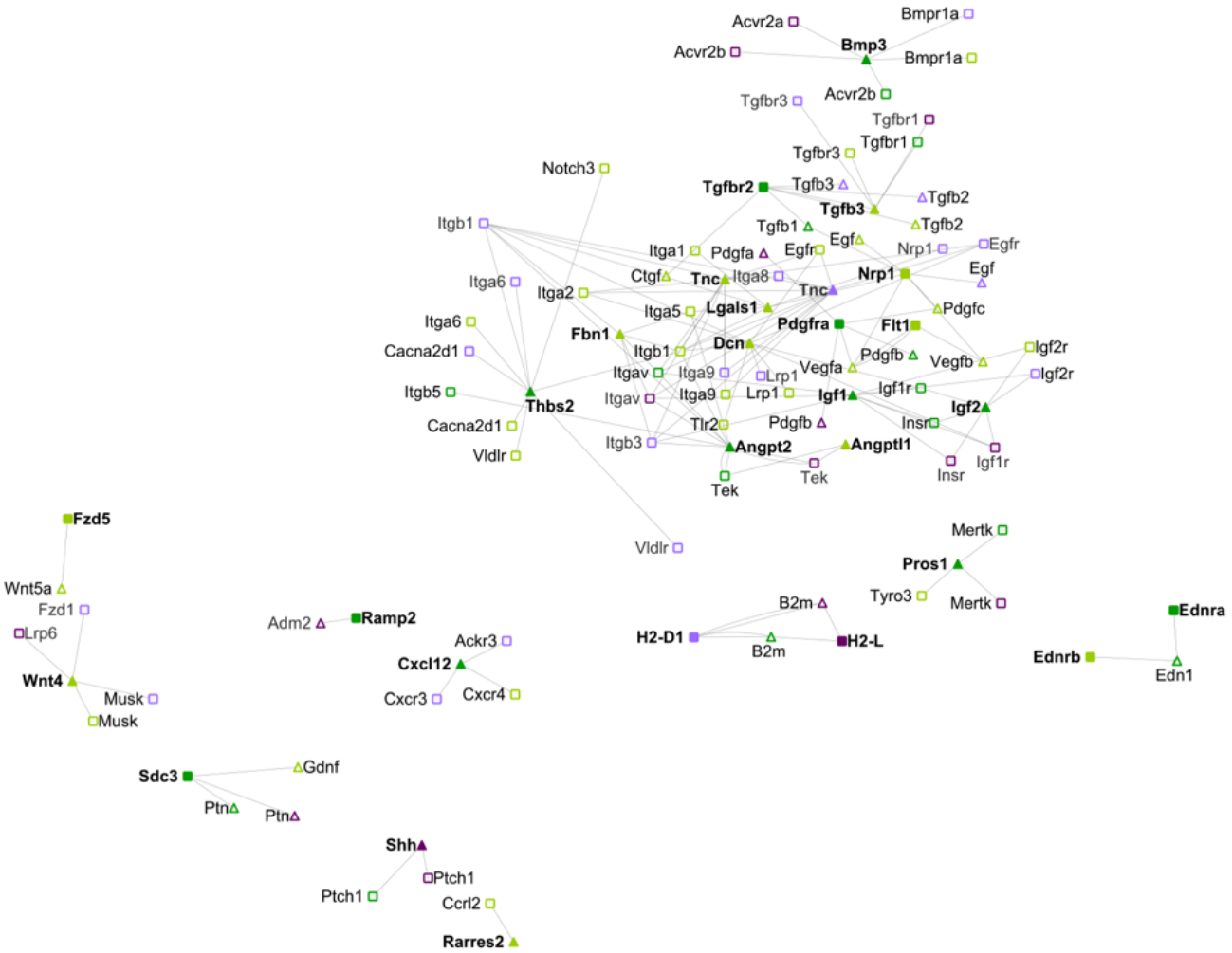
Intercellular networks between oocytes and somatic cells from primary to secondary transition. Legend as in **Fig. 1**. The connections and the references for each of them can be found in **File S1**.

**Figure S2.**
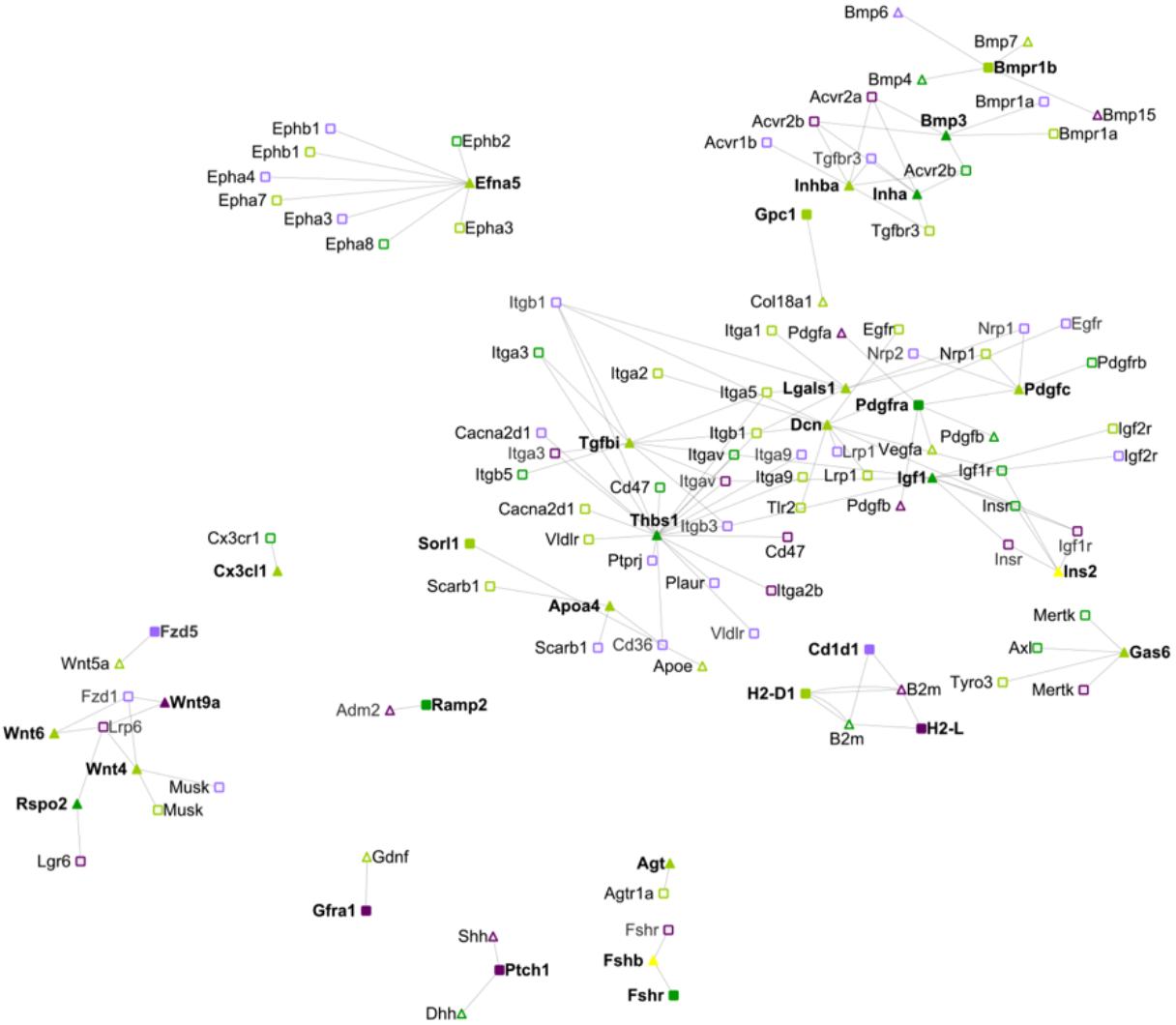
Intercellular networks between oocytes and somatic cells from secondary to small antral transition. Legend as in **Fig. 1**. The connections and the references for each of them can be found in **File S1**.

**Figure S3.**
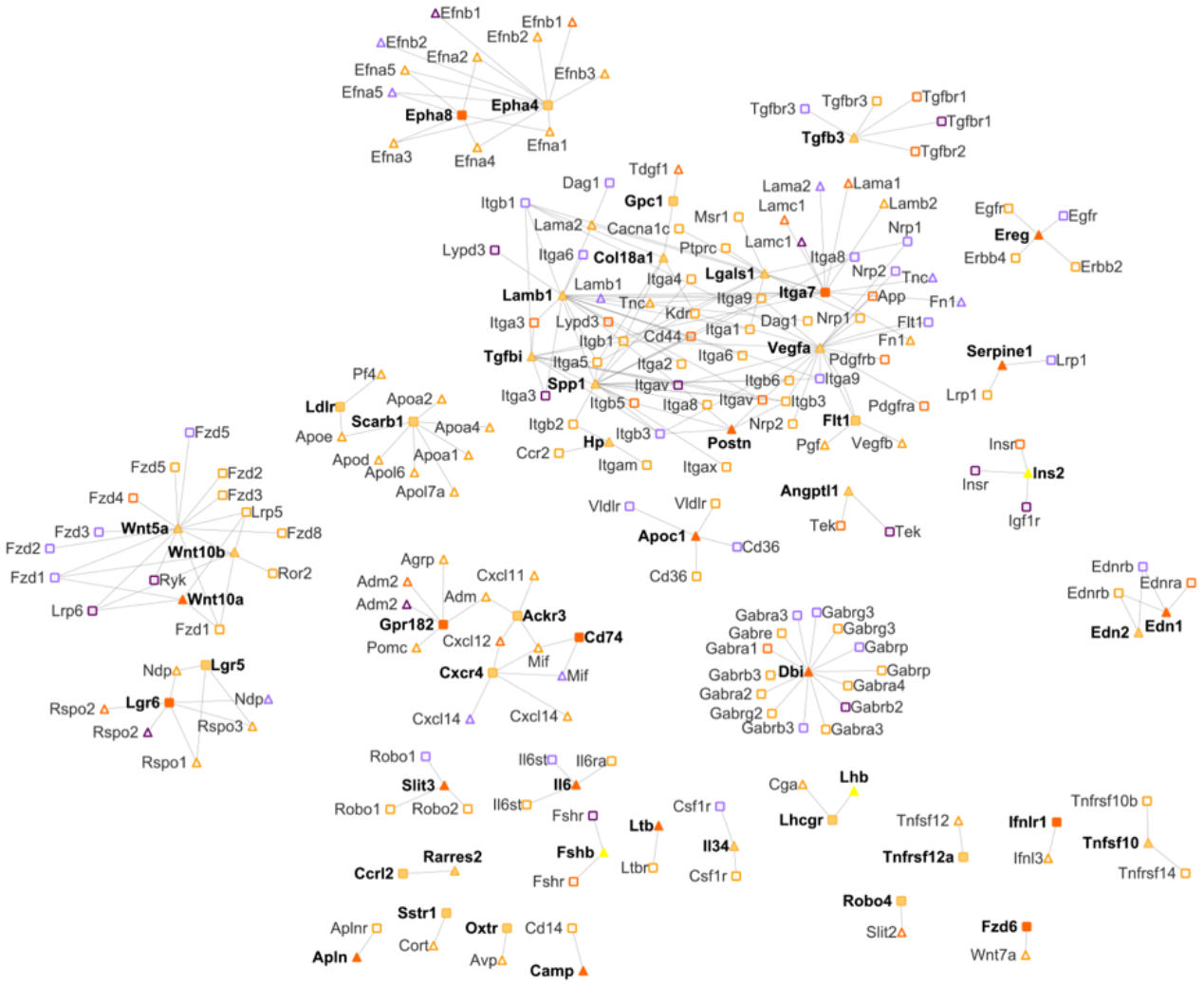
Intercellular networks between oocytes and cumulus cells from small antral to large antral transition during the NSN to SN transition. Legend as in **Fig. 2**. Red, cumulus cells; blue, mural cells; yellow ligands from the endocrine system; orange, cumulus cells during the NSN to SN transition. The connections and the references for each of them can be found in **File S1**. Oocyte: FC≥2.5, p-value≤0.01; transition cumulus granulosa cells: FC≥1.5 and p-value≤0.01

**Figure S4.**
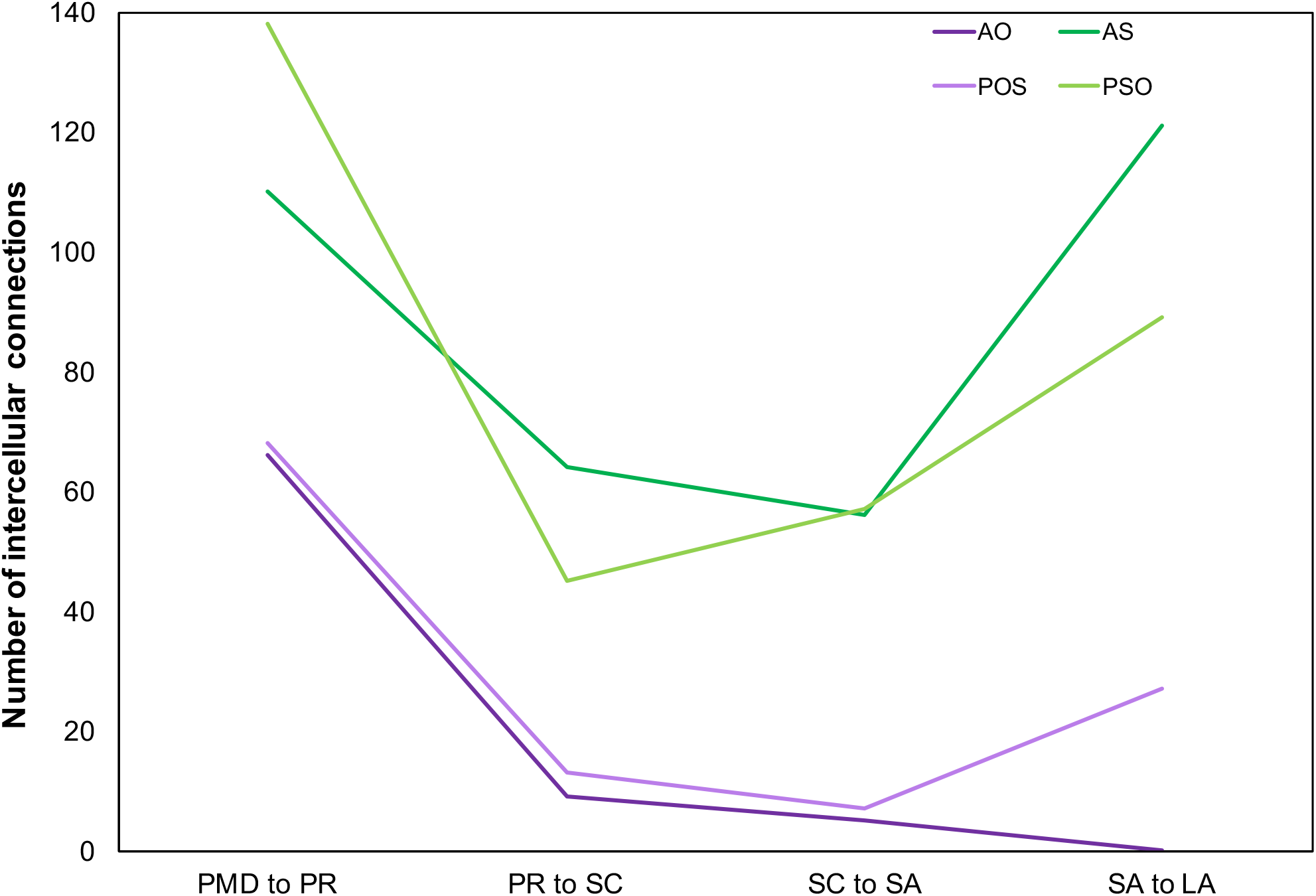
Temporal evolution of the number of autocrine and paracrine connection during ovarian follicle development. PMD, primordial; PR, primary; SC, secondary; SA, small antral; LA, large antral; AO, oocyte autocrine communications; AS, somatic autocrine communications; PO, paracrine communications between oocyte ligands and somatic receptors; PS, paracrine communications between somatic ligands and oocyte receptors.

**Table S1.**
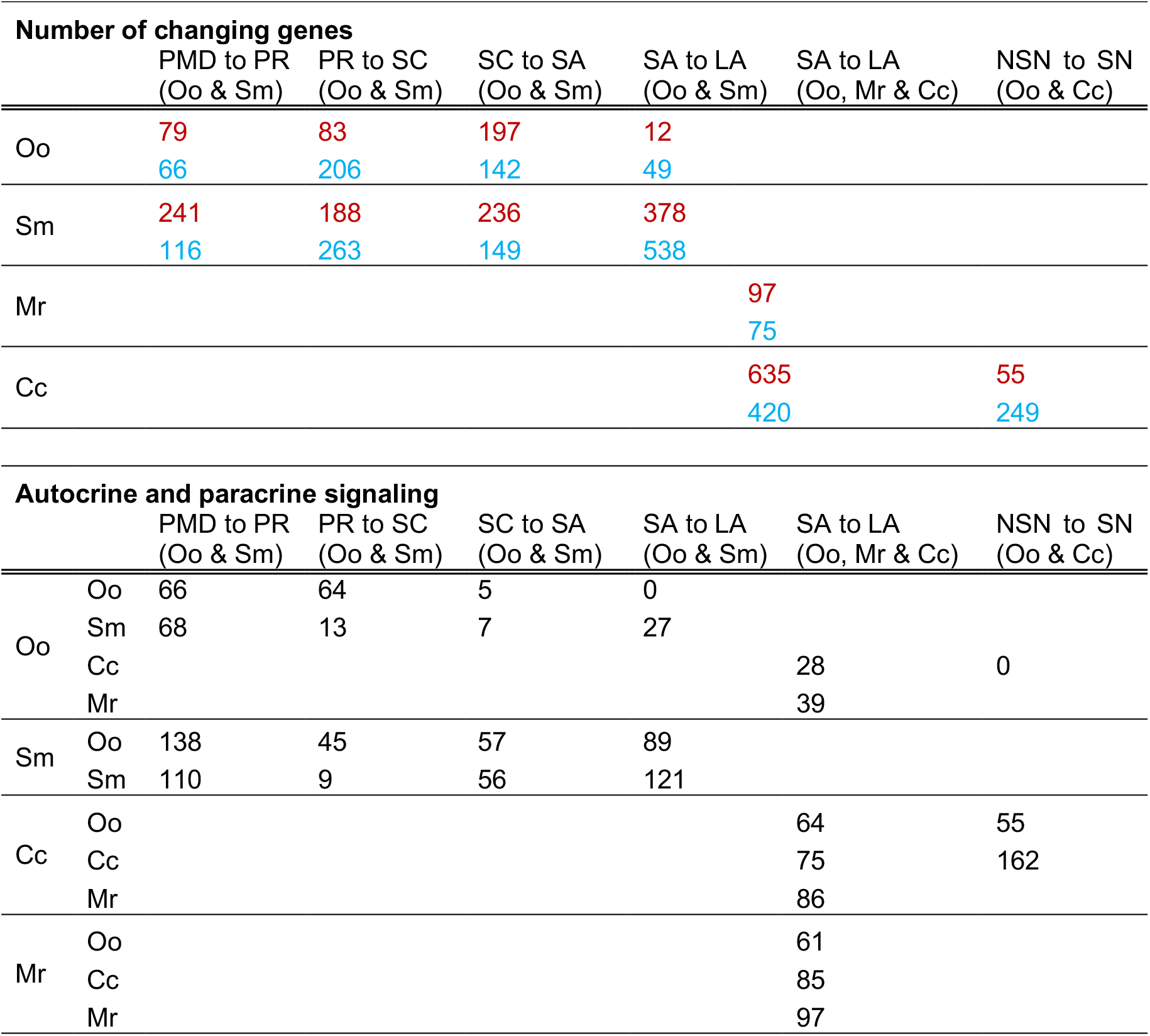
Number of upregulated (blue) and downregulated (red) transcripts during ovarian follicle development and the number of paracrine and autocrine communications between the different ovarian cell types. PR, primary; SC, secondary; SA, small antral; LA, large antral; NSN, antral follicle that present an oocyte that has a non-surrounded nucleolus; SN, antral follicle that present an oocyte that has a surrounded nucleolus; Oo, oocyte; Sm, somatic cells (i.e., granulosa and theca cells); Mr, mural cells; Cc, cumulus cells.

## SUPPLEMENTARY INFORMATION

### Supplemental Note 1

During the primordial to the primary ovarian follicle transition, the oocyte starts growing from 20 to 40 μm on average, forming the zona pellucida around the oocyte(*66*), which is dictated by the presence of the zone pellucida proteins (i.e., ZP1, ZP2 and ZP3), and gap connections (e.g., connexin-37 (*67*) and connexin-43 (*68*)). The transcripts of several zona pellucida proteins were present only in the oocyte *Zp1*, *Zp2* and *Zp3* (**File S2**). The major component of the gap connections during ovarian follicle development is connexin-37 (*67*) between the oocyte to granulosa cells and connexin-43 (*68*) between granulosa cells and both were greatly increased in the oocyte and somatic cells, respectively (**File S2**).

Granulosa cells starts changing their phenotype, from cuboidal cells to squamous cells and the formation of the basal lamina by deposition of extracellular components around the granulosa cells, collagen alpha-1, 2, alpha-3, alpha-4 and alpha-5 (*69*). While the transcripts that encode for collagen alpha-1 and alpha-2 were presented in the somatic cells and in the oocyte, they were downregulated in the somatic cells with no significant change in the transcripts abundance in the oocyte (**File S2**). Interestingly, the transcriptional abundances of laminin alpha-2 and laminin beta-1 transcripts was increased (**File S2**).

The inhibition that maintained the primordial follicle in this quiescent stage is therefore removed so that the follicle can transition into the primary stage. Amhr2, the gene that encodes for the receptor AMHR, increased its abundance in the somatic cells (**File S2**) so that the somatic cells began to be sensitive to endocrinal changes such as FSH (*70*). The transcript that encodes for the FSH receptor, *Fshr* increased during the primordial to primary transition, although it was not significant in somatic cells (**File S2**). In fact, the transcriptional level of *Fshr* commenced to be above the background level of the transcriptional arrays in 4-day old mice (p-value<0.01).

### Supplemental Note 2

During the primary to secondary follicles transition, the follicle start acquiring up to 10 layers of granulosa cells (*35*), the formation of the theca layer commences and the follicles are capable to produce estrogenic hormones. Ovarian follicle development growth factor *Gdf9* transcript was highly abundant in the oocyte(*71*) and its expression level was similar to that in primary follicles (**File S2**). *Igf1* transcriptional abundance during this period changed(*72*), but only in somatic cells. Several theca cell markers were presented in the ensemble transcriptomic data during the primary to secondary transitional period(*42*), such as *Cyp17a1* whose abundance was increased only in somatic cells—it was undetected in the oocyte transcriptome. Similarly, *Cyp11a1* abundance increased during this period in both the oocyte and the somatic cells (**File S2**). Interestingly *Cyp11a1* has been mostly studied in terms of somatic cells, although it has been reported to be present in the oocyte of zebrafish (*73*).

### Supplementary Note 3

A few secondary follicles more sensitive to the endocrinal hormones FSH and LH mature into antral follicles and the rest succumb into atresia, the default pathway (*37*). Follicles during this transition start producing androgens in the theca cells and estrogens in the granulosa cells (*38*) and the antrum cavity filled with hyaluronic acid and proteoglycans(*39, 40*)-versican and perlican (*41*) emerges. Additionally, theca cells become vascularized (*42*). The receptors *Fshr* and *Lhr* transcriptional abundance slightly increased in the somatic cells (**File S2**). Surprisingly, most of the genes that encode for commonly known androgen producing enzymes were not detected (i.e., *Star*) or did not significantly changed their transcriptional levels in this transition (i.e., *Cypa11, Cyp17a1, Hsd3b1*), while genes that encode for estrogenic enzymes significantly increased, e.g., *Cypa19a1* and *Hsd17b1* (**File S2**). Interestingly, *Cypa19a1* was also significantly increased in the oocyte (**File S2**), which agrees with observations that indicated that *Cypa19a1* is present in isolated human oocytes (*74*). Similarly, angiogenesis ligands *Vegfa* and *Vegfb* and the *Vegfgr-1* receptor were above the background of the transcriptional results, yet their abundance was not significantly altered during the secondary to small antral transition. Genes that encode for enzymes implicated in ovarian follicle vascularization had opposite transcriptional behaviors. For instance, while *Angtp1* abundance was not detected in somatic cells, *Angtp2* transcriptional abundance significantly increased (**File S2**). *Angtp2’s* receptor, *Tek*, was ubiquitously present in all the stages during ovarian follicle development in somatic cells and since secondary follicles onwards on the oocyte. Versican (*Vcan*) transcript was found only in somatic cells since very early stages in follicle development and perlican (*Hspg2*) was also present during ovarian follicle development, been initially observed in primordial follicles. However, transcripts encoding for hyaluronic acid (i.e., *Has1, Has2, Has3*) were not detected during this transition, although *Has3* was detected in prior transitions.

### Supplemental Note 4

At the end of the transition between the small antral to large antral, the oocyte is competent to resume meiosis (*43*), the granulosa cells have differentiated into mural and cumulus granulosa cells and the antrum cavity is fully formed. At the transcriptional level, *Fshr* abundance did not significantly changed at this stage, while *Lhcgr* transcriptional abundance was highly increased in the somatic cells. Most of the commonly known enzymes that catalyze the production of androgens in the theca cells were now highly changed(*42*), e.g., *Star, Cyp11a1, Cyp17a1* and *Hsd3b1*. Similarly, estrogenic enzymes well known to be present in the granulosa cells, their genes were significantly changing, such *Cyp19a1*—note that *Cyp19a1* transcript was not significantly activated in the oocyte anymore. Only *Hsd17b1* did not significantly changed its abundance. Transcriptomics data also confirmed that that angiogenic factors seemed to play an important role during the large antral formation, e.g., *Flt1* gene that encode for VEGF-A (*Flt1* gene) and its corresponding receptor, Vegfr-1 significantly increased their abundance during antrum formation (**File S2**). On the other hand, while *Angpt1* increased its transcriptional level in somatic cells, *Angtp2* abundance had the opposite effect. Two of the major proteogleocans that have been identified in the follicular fluid, versican (*Vcan*) and perlican (*Hspg2*), were significantly changed at that transcriptional level during the antral formation. Taken it together, the ensemble of transcriptional available data confirmed the well-known small to large antral transition.

### Supplemental Note 5

At the end of the small to large antral ovarian follicle transition, the granulosa cells are fully differentiated into mural granulosa cells and the cumulus granulosa cells. At the transcriptional level, cumulus and mural granulosa cells continued producing and increasing the levels of *Fshr*, yet only mural cells abruptly increased the transcriptional of *Lhcgr*—confirming prior observations at this stage (*75*). Most of the genes that encode for estrogenic enzymes had significantly increased their transcript abundance in mural cells (e.g., *Star, Cypa11*) and only a few were also in both mural and cumulus cells (e.g., *Cyp19a1*). Other estrogen enzymes transcripts were still being produced yet none of them were significantly changing during the small to large antral formation, neither in the cumulus nor in the mural granulosa cells (e.g., *Hsd3b1, Hsd17b*). Interestingly, *Cyp17a1*—the gene that encode for the enzyme CYP17A1 and is key for the production of testosterone—was neither detected in the cumulus nor in the mural granulosa cells, which maybe an indication that CYP17A1 is theca cell specific enzyme. Other transcripts have been reported in the literature as mural markers, such as *Cd34* (*76*), which was indeed only present in the mural cells, yet *Slc38a3*—another mural granulosa cell marker—was significantly decreased during the antral formation, not only in mural but also in the cumulus granulosa cells. Moreover, *Scl38a3* mural transcription was below the detection limit in the large antral follicles, while it was still highly transcribed in the cumulus cells. In the case of cumulus granulosa cell markers, such as *Amh*, most of them were either not present in any of the granulosa cell types or were non-significant changed during the transition (e.g., *Ar*) not only by cumulus cells but also by mural (**File S2**), which contradict prior research results (*76*).

In terms of angiogenesis markers, neither the cumulus not the mural granulosa were involved in the process, according to our results. For instance, *Vegfb* transcript was not significantly increased during this stage neither in cumulus or mural granulosa cells. The transcripts of *Vegfa* and its receptor *Vegfr-1* were not detected neither in cumulus not mural cells, which could be an indication that they are only transcribed in theca cells, and thus being theca cell-specific genes. Similar transcriptional behavior was observed in for *Angpt1* and *Angpt2* (**File S2**).

Finally, components of the extracellular matrix such as *Vcan* were significantly increasing their transcriptional rates at this stage but only in mural cells. On the contrary, *Hspg2* was not transcribed neither by the cumulus cells not the mural cells, which agrees with prior observations that perlican is produced by theca cells (*77*). Transcripts for hyaluronic-producing enzymes HAS1 and HAS3 were non-detected neither in cumulus or mural cells and only cumulus cells significantly increased the production of the *Has2* transcript (**File S2**).

### Supplemental Note 6

Oocyte maturation is required for adequate egg fertilization (*44*) and entails the nuclear maturation of the oocyte: chromatin condensation in the oocyte nucleolus, from a non-surrounded nucleolus (NSN) to a surrounded nucleolus (SN) (*45*). This transition cannot be achieved by nude oocytes; they required the present of cumulus granulosa cells(*46*).

During the NSN to SN transition, we identified a total of 249 genes that were upregulated, while only 55 were downregulated in cumulus cells (**Table S1**). Cumulus specific genes during the oocyte nuclear maturation were significantly changing. For instance, *Hbb-b2* (hemoglobin beta adult minor chain) was the most downregulated gene in the system, followed by *Nrsn1* (neurensin 1) and *Smim5* (small integral membrane protein 5). Yet, *Bhmt* (betaine-homocysteine methyltransferase), *Pik3c2g* (phosphatidylinositol-4-phosphate 3-kinase catalytic subunit type 2 gamma) and *Apln* (apelin) were the top three most upregulated genes in the system (**File S2**).

## SUPPLEMENTARY FILES

**File S1**. Inter-cellular connections *in vivo* during ovarian murine follicle development

**File S2**. Transcriptional abundance of selected genes in the oocyte, somatic, cumulus and mural cells during ovarian follicle development. This file has been adapted from **File S8** from Peñalver Bernabé and colleagues (*17*)

**File S3**. The most likely involved TFs during ovarian follicle development.

